# Error encoding in human speech motor cortex

**DOI:** 10.1101/2025.06.07.658426

**Authors:** Xianda Hou, Carrina Iacobacci, Nicholas S. Card, Maitreyee Wairagkar, Tyler Singer-Clark, Erin M. Kunz, Chaofei Fan, Foram Kamdar, Nick Hahn, Leigh R. Hochberg, Jaimie M. Henderson, Francis R. Willett, David M. Brandman, Sergey D. Stavisky

## Abstract

Humans monitor their actions, including detecting errors during speech production. This self-monitoring capability also enables speech neuroprosthesis users to recognize mistakes in decoded output upon receiving visual or auditory feedback. However, it remains unknown whether neural activity related to error detection is present in the speech motor cortex. In this study, we demonstrate the existence of neural error signals in speech motor cortex firing rates during intracortical brain-to-text speech neuroprosthesis use. This activity could be decoded to enable the neuroprosthesis to identify its own errors with up to 86% accuracy. Additionally, we observed distinct neural patterns associated with specific types of mistakes, such as phonemic or semantic differences between the person’s intended and displayed words. These findings reveal how feedback errors are represented within the speech motor cortex, and suggest strategies for leveraging these additional cognitive signals to improve neuroprostheses.

## Introduction

Humans and other animals can monitor the outcomes of their actions. Identifying errors through sensory feedback is a crucial process that helps refine actions to improve future performance^1^. This ability underlies a broad range of learning processes and is observed in behavioral phenomena such as post-error slowing^2,3^ and within-trial adaptation during motor tasks^4,5^. Understanding an integrative behavior like sensorimotor learning requires identifying where in the brain error-related activity occurs.

The neural mechanisms supporting error detection and correction have been characterized using multiple measurement modalities, including EEG^6,7^, fMRI^8^, and intracranial recordings^9^. Growing evidence suggests that error-related signals are observed in a distributed network of cortical and subcortical regions, extending beyond the traditionally emphasized roles of the medial frontal cortex and dopaminergic system^6,8,7,10^. These feedback-related error signals reflect mismatches between predicted and actual outcomes, providing the basis for cognitive and motor-level adjustments^11,12,8,13–15^. For example, studies in animal models and computational simulations have shown that motor cortex activity can change on a trial-by-trial basis in response to feedback about reaching behavior, supporting flexible and efficient learning^16,5,17,1^. In the context of brain–computer interfaces (BCIs), where errors arise from the neural decoder’s misinterpretation of the user’s intended actions, outcome-related error signals have been detected in dorsal (hand) motor cortex during cursor control tasks^18–20^, demonstrating that neural populations directly involved in motor execution can also encode information about task success or failure.

Here we asked the question: are similar error signals observed in the human speech motor cortex, a region that supports a distinct and complex motor behavior? Successful production of speech requires precise control of articulator movements, and it also conveys rich semantic and linguistic meaning. Subtle perturbations in speech output, such as shifts in formant frequencies or voice onset times, can disrupt not only its acoustic quality but also its perceived linguistic content^21^. Despite this complexity, speakers can adapt their vocal output in real time, maintaining intelligibility even when feedback is distorted^22,23^. However, the neural basis of this adaptive capacity—and whether it involves an internal monitoring signal reaching the motor cortex akin to what has been found in forelimb and BCI motor adaptation—remains poorly understood, especially at the resolution of neuronal action potentials.

Understanding how the speech motor cortex processes and adapts to feedback has both neurobiological and translational significance. From a scientific perspective, it offers a window into how the brain maintains its capacity to verbally transmit communicative intent with high fidelity under dynamic and noisy conditions. From an engineering perspective, understanding speech error encoding directly informs the development of better BCIs, which are an emerging class of medical devices that have recently shown promise in restoring communication for individuals with severe speech impairments^24,25^ by decoding neural signals into text (“brain-to-text”)^26–28^. Despite recent advances in speech BCI decoding accuracy, current systems remain prone to decoding errors and rely heavily on extensive labeled training data—either from prompted speech, or self-generated speech that the user retrospectively labels as correct^28^—for updating the decoder. These errors often stem from similar neural representations of acoustically or motorically similar words^26–28^ and can significantly slow communication by requiring user corrections. Leveraging intrinsic neural error signals offers a potential solution: if such signals could be detected from the same electrode arrays already in place for speech decoding and identified on a single-trial basis in real-time, the BCI could assess its own performance and automatically initiate corrective changes without requiring the user’s explicit feedback.

In this study, we show that speech task outcome error signals are present in the human ventral precentral gyrus (speech motor cortex) and can be reliably decoded (90% accuracy during open-loop word copy task and 86% during a speech BCI task) in real time. Beyond detecting whether an error occurred, the neural activity also encodes higher-level linguistic differences between incorrect feedback and the expected word. These signals exist in the same cortical locations targeted with electrode arrays that enable a high-performance speech neuroprosthesis. Finally, we demonstrate a prototype online error detection system capable of accurately predicting the correctness of a word decoded by the speech BCI within 550 ms of it being presented to the user.

## Results

To investigate the presence of neural error signals in the human speech motor cortex during speech BCI use, we analyzed multielectrode array recordings from BrainGate2 clinical trial participants T15 and T12 while they attempted speech tasks. At the time of data collection, both participants were severely dysarthric due to amyotrophic lateral sclerosis (ALS). Participant T15 had four arrays implanted along his ventral precentral gyrus (vPCG) spanning the functional regions of area 6v, 4 and 55b; array locations are depicted in **Fig. 1b**. Participant T12 had two arrays implanted in the area 6v of vPCG and 2 arrays in area 44 of the inferior frontal gyrus (IFG); array locations are shown in **Supplemental Fig. 2**.

**Fig. 1.**
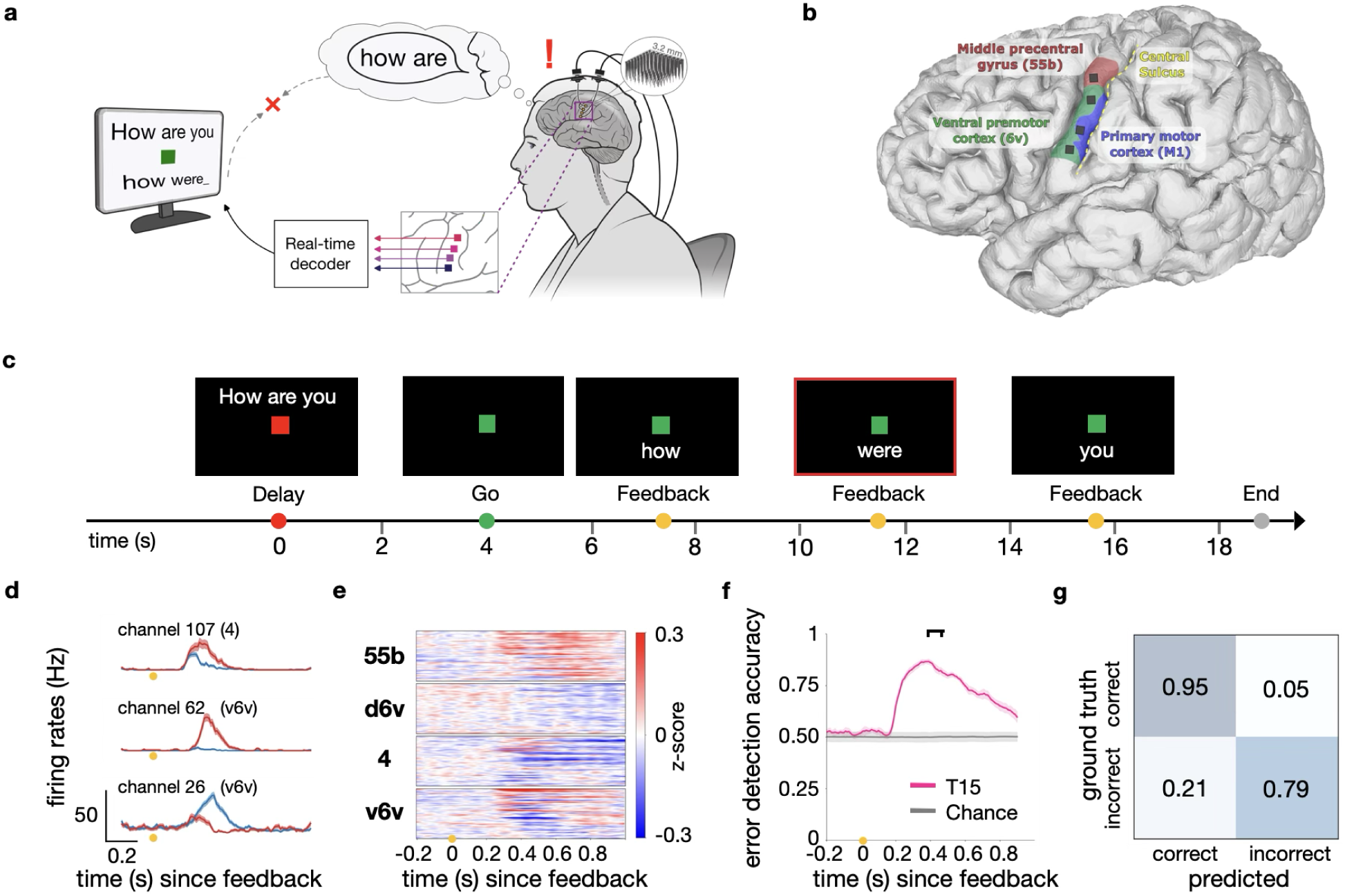
Neural error signals during a brain-to-text task with sequential feedback. **a**) Schematic of the brain-to-text speech BCI. Neural activity recorded from four microelectrode arrays was processed and decoded into words. **b**) Approximate microelectrode array locations (colored squares) superimposed on a 3-D reconstruction of participant T15’s brain. Colored regions correspond to cortical areas aligned to the participant’s brain using the Human Connectome Project MRI protocol scans before implantation^29^. **c**) Timeline showing an example trial of the Sequential Feedback Task. The participant was shown the prompted sentence during a delay period, after which he attempted to speak that sentence. Each screen update illustration highlights how only the most recently decoded word was displayed.In this figure (not in the actual task), the frame containing an incorrectly decoded word is highlighted with a red border. **d**) Selected electrodes’ trial-averaged multi-unit firing rates (mean ± s.e.) aligned to correct (blue) and incorrect (red) feedback display (n=335 trials; 215 correct and 120 incorrect). **e**) Normalized trial-averaged spike band power difference between trials in which the feedback (decoded word appearing on-screen) showed the incorrect versus correct word. The vertical axis shows 256 electrodes which are grouped by array. **f**) Offline classification accuracy of the correctness of the feedback (mean ± s.d.) as a function of the start of a 100-ms sliding window of neural data. **g**) Confusion matrix of the classification results using the window [390, 490] ms (black bracket in f) after feedback display.

We designed both open-loop and closed-loop speech tasks to investigate neural error signals induced by visual and auditory feedback following the participant’s speech attempts. In the closed-loop paradigm, the BCI predicted the attempted word from neural activity, updating dynamically as each trial unfolded (**Fig. 1a**; see^28^ for further details). In contrast, the open-loop tasks presented feedback drawn from a predefined set of possible words, exploring how different types of incorrect words influenced neural responses.

### Neural error signals can be decoded from the human speech motor cortex

To assess neural responses to erroneous speech feedback, we first analyzed data from participant T15 using a brain-to-text BCI with sequential feedback. Unlike standard brain-to-text tasks^27,28^, where the decoded sentence continuously updates by building out word-by-word on the screen, the “Sequential Feedback Task” displays only the most recent predicted word at a fixed location on a screen (**Fig. 1c**). This task design minimizes eye movement and focuses the participant’s attention on discrepancies between the intended and decoded words.

Neural activity showed significant error-related modulation beginning approximately 250 ms after visual feedback, with strong error-related responses measured from the electrode arrays located in areas v6v, 4, and 55b of the precentral gyrus (**Fig. 1e**). Multiunit activity exhibited a variety of responses, with some increasing firing rates after erroneous feedback and others decreased firing (**Fig. 1e**). Using a 100-ms sliding window, a Support Vector Machine (SVM) classifier could predict feedback correctness significantly above chance starting at 200 ms, peaking at 86.6% accuracy at 390 ms post-feedback (**Fig. 1f**), with a false positive rate of 5% (**Fig. 1g**). The arrays that most contributed to speech decoding also encoded error-related information (**Supplemental Fig. 1**); the array in d6v did not significantly contribute to error detection and was excluded from subsequent analyses.

While the aforementioned brain-to-text speech BCI task is a relatively naturalistic communication behavior, a downside of that task is that there were a number of rapidly occurring task events competing for the participant’s attention and a high degree of variability in the words being spoken. To more systematically study these neural error signals, we had T15 perform an open-loop ‘Word Copy Task’ and we also replicated this experimental paradigm with participant T12, who had two arrays in vPCG. Each trial prompted the participant to try to speak a single word, after which they were provided with one of five types of feedback: 1) the cue word (i.e., correct feedback), 2) a pseudoword differing from the attempted word by one phoneme, 3) a homophone, 4) a minimal pair real word (i.e., differing from the attempted word by one phoneme), or 5) a synonym (**Table 1** lists all cue-feedback pairs).

**Table 1.** Cue-feedback conditions used in the Word Copy Task.

| Cue | Pseudoword | Homophone | Minimal pair | Synonym (BERT dist) |
| --- | --- | --- | --- | --- |
| feet | reet | feat | meet | paws (0.256) |
| son | sen | sun | sin | child (0.264) |
| fore | fure | for | fare | front (0.187) |
| dear | deat | deer | dean | beloved (0.263) |
| heal | heam | heel | heat | cure (0.119) |
| vain | tain | vein | pain | useless (0.155) |

For both participants, neural responses in speech motor cortex arrays exhibited clear tuning to feedback correctness (**Fig. 2c,d, Supplemental Fig. 2d,h**). Significant error-related modulation was observed in both participants starting 250 ms after feedback, with the highest classification accuracy of 90.8% (T15) and 77.9% (T12) at 460 ms and 430 ms after feedback display (**Fig. 2e**). T15 demonstrated stronger neural modulation, likely due to having more arrays in precentral gyrus as well as better array recording quality. Unlike T15, T12 also had two Utah arrays in her inferior frontal gyrus (IFG). Neural activity recorded from these IFG arrays also exhibited strong error-related modulation (**Supplemental Fig. 2c,g**), supporting prior evidence of IFG involvement in conflict resolution during sentence parsing^30,31^.

**Fig. 2.**
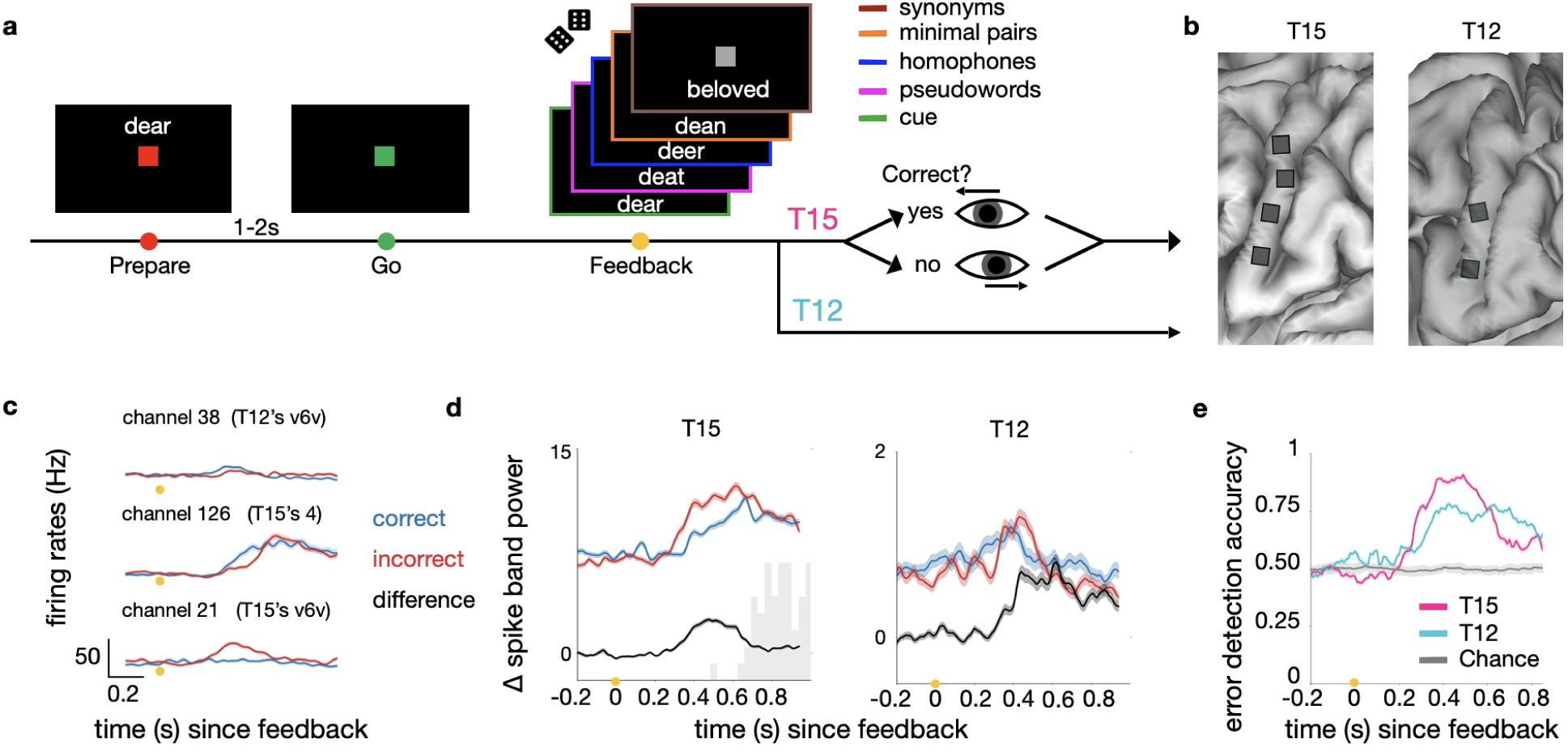
Neural error signals during a Word Copy Task. **a**) Illustration of the Word Copy Task. Participant T12’s trials ended two seconds post-feedback, whereas participant T15 additionally was prompted to confirm feedback correctness using an eye tracker. **b**) Approximate placement of microelectrode arrays (gray squares) on the vPCG of both participants. **c**) Trial-averaged firing rates from selected electrodes showing neural responses to correct (blue) versus incorrect (red) feedback (for T15, n=161 trials; 68 correct and 93 incorrect; for T12, n=243 trials; 125 correct and 118 incorrect). **d**) Neural population modulation (mean ± s.e.) in spike band power (z-scored) across all vPCG arrays. Red and blue lines show neural distances between incorrect and correct feedback responses versus a “do nothing” baseline condition. The black line shows the neural distance between incorrect and correct feedback responses. Note the different vertical scales between participants; T15 had more electrodes and they recorded stronger speech-related modulation. The light gray histogram (left panel) indicates the time distribution of when T15 initiated a response with a saccade. **e**) Offline classification accuracy (mean ± s.d., leave-one-cue-out cross validation) of feedback correctness as a function of the start of a 150-ms sliding window.

### Encoding of language-related information in error signals

Beyond detecting a binary outcome error signal in speech motor cortex, we further explored whether the neural responses following feedback also encoded the linguistic relationships between the attempted and feedback words. Both participants had electrodes with distinct tuning patterns to different error types (**Fig. 3a**). Population-level analyses further revealed distinctive modulation patterns in response to the different kinds of incorrect feedback responses (**Fig. 3b**, **Supplemental Fig. 3a&d**). To verify that these neural differences did not just result from low-level differences in reading the specific words (e.g., phonemic tuning), we tested if a classifier trained on a subset of cue-feedback word sets could predict error feedback types for unseen word sets. For T15, the classifier successfully differentiated pseudowords, homophones, and synonyms with an overall 59.4% accuracy (well above chance) at 410 ms post-feedback (**Fig. 3c,d)**, although minimal pairs were not identified above chance. Although T12’s overall accuracy with her vPCG arrays did not surpass chance, certain arrays achieved above-chance decoding for specific feedback types (**Supplemental Fig. 3)**. We attribute this poor ability to decode between types of error signal (in contrast to above-chance accuracy for the easier binary classification of whether the word was correct or not) to the much lower signal-to-noise ratio in T12’s vPCG arrays. Interestingly, T12’s IFG arrays, which had higher signal quality, demonstrated strong language-related error type classification performance (**Supplementary Fig. 4**).

**Fig. 3.**
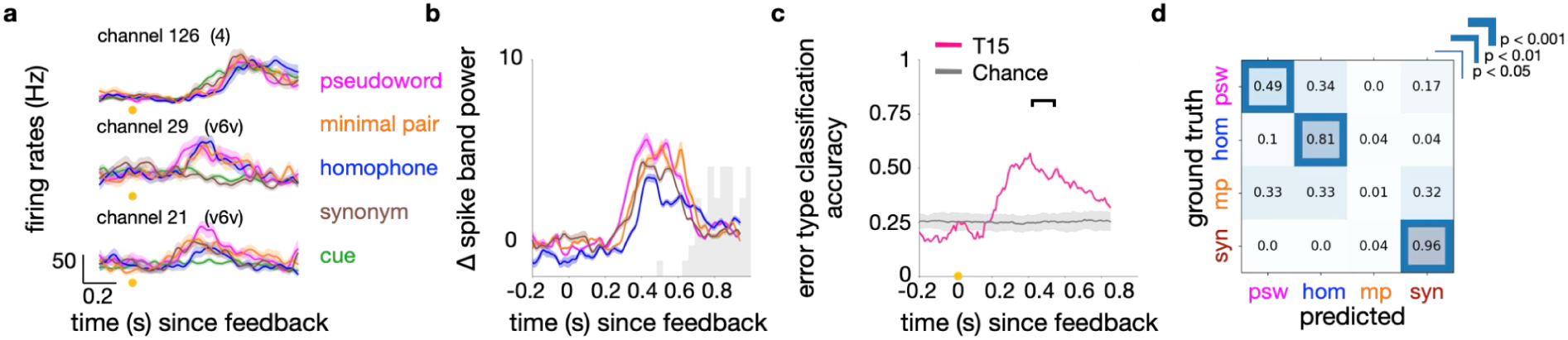
Neural error signals encode not only correctness but also error type. **a**) Selected electrodes’ trial-averaged firing rates, showing neural responses around the presentation of different types of incorrect feedback (n=93 trials, pseudoword:minimal pair:homophone:synonym=24:23:22:24 trials). **b**) Neural population modulation (mean ± s.e.) in spike band power for participant T15. Modulation was quantified as the neural distance between ensemble activity following correct feedback and following each type of incorrect feedback. The light gray histogram indicates the time distribution of when participant T15 initiated a response with a saccade. **c**) Offline classification accuracy (mean ± s.d.) of error type using a 150 ms sliding window of neural data across time. **d**) Confusion matrix showing classification performance at 410 ms after feedback display, highlighting the discriminability across incorrect feedback types. Significant diagonal cells (Shuffle test with 1000 times of label permutations, Bonferroni-corrected) are highlighted with colored boxes of varying thickness.

### Online error detection during real-time speech BCI use

Having found a reliable neural error signal following erroneous text feedback, we investigated the feasibility of adding automatic neurally triggered error detection to participant T15’s brain-to-text speech neuroprosthesis. We started by first analyzing previously collected data where T15 performed a standard speech BCI sentence-copy task^28^. In this task, he was prompted to speak a full sentence following a cue on the screen. As he attempted to speak, the predicted partial sentence was displayed and updated while an eye tracker measured where he was looking on-screen. We measured the neural activity in the time epochs when the participant looked at each feedback word (see Methods). Consistent from findings from the Sequential Feedback Task described earlier, we also observed error-related modulation in v6v, 4 and 55b (**Fig. 4b**). However, although the error detection accuracy was still above chance (65.7% ± 0.33., p=3.02e-13), it was substantially lower than in the Sequential Feedback or Word Copy Task. This could reflect the limitations of analyzing data from this more complex task (e.g., there’s no guarantee that the participant read a given word exactly when he fixated near it^32,33^ and he may have been more focused on articulating the upcoming words). In addition, the participant may have attended less to individual word errors because the neuroprosthesis’ language model would correct many mistakes using context-dependent predictions as the sentence unfolded, reducing the perceived importance of early mistakes. Thus, the attenuated error signals we observed in this standard speech BCI task could be explained by the theory that the generation of error-related negativity is modulated by the subjective significance of the error to the individual^3^.

**Fig. 4.**
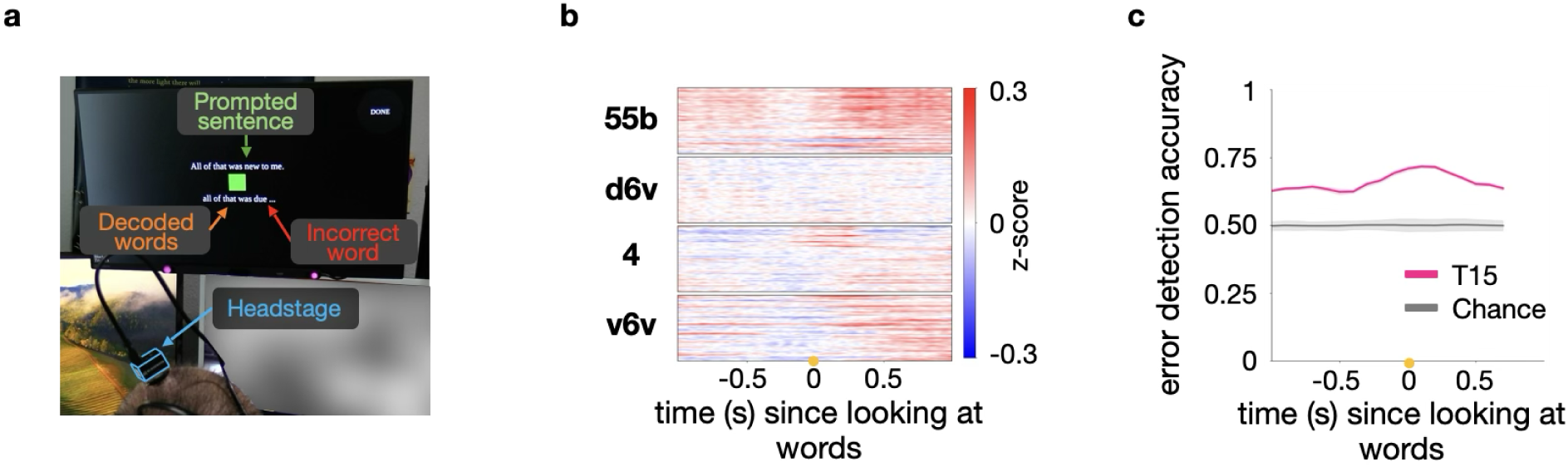
Neural error signals during naturalistic speech BCI use. **a**) Participant T15 performing the Sentence Copy Task using the speech neuroprosthesis. T15’s attempted words were decoded using the pipeline in Fig. 1a. **b**) Normalized trial-averaged spike band power difference after the participant looked at incorrect words compared to after he looked at correct words (n=1048 trials; 532 correct and 516 incorrect). The vertical axis shows 256 electrodes which are grouped by array. **c**) Offline classification accuracy (mean ± s.d.) when detecting whether a displayed word was correct or not from the neural data. Accuracy is shown as a function of the start of a 100-ms sliding window aligned to when the participant moved his gaze onto the word.

In light of the previous results, we predicted that a task structure that encouraged focused engagement by the participant on the decoded words as they were presented would help elicit stronger error-related neural modulation. Thus, to demonstrate a proof-of-principle error detecting speech BCI, we modified the speech neuroprosthesis user interface to provide feedback less frequently and emphasize each word as it was presented. In this Emphasized Feedback Task, the participant could still see the history of previously decoded words, but in smaller font, whereas the latest decoded word was highlighted prominently (**Fig. 5a**). The participant was instructed to pay attention to the latest decoded word, which would normally show up within 100 milliseconds after his speech attempt.

**Fig. 5.**
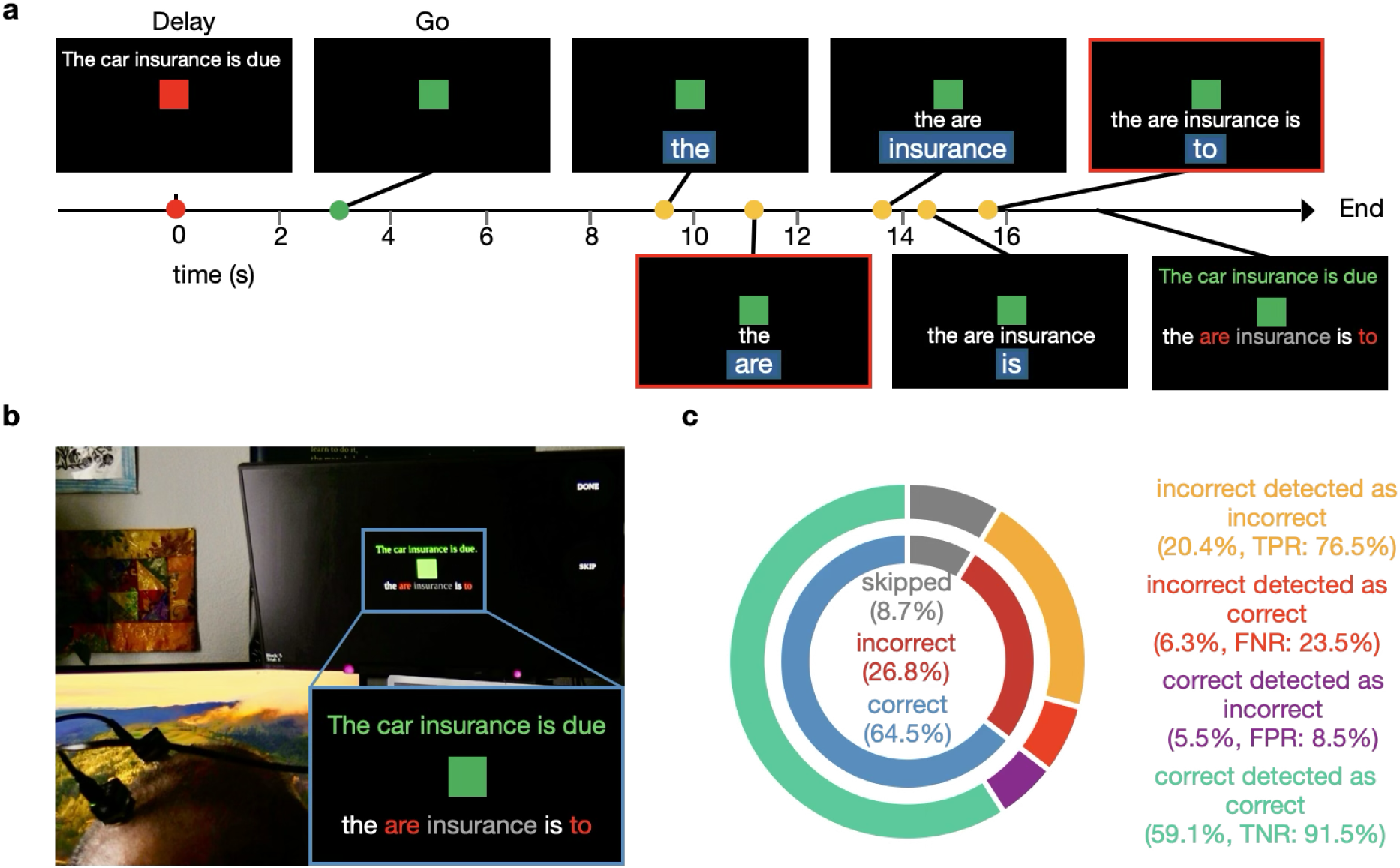
Online neural error signal detection in a speech BCI. **a**) Example trial structure during BCI use with emphasized feedback. The timeline shows how the screen updated as the participant attempted to say the target sentence that was prompted during the delay period. The most recently decoded word is highlighted with a blue background and was subsequently appended to the previously decoded words, which are shown below the green square. **b**) Photograph of the participant looking at the finalized sentence. Words identified by the neural error decoder as incorrect are highlighted in red, while words in gray represent “skipped words”: those with display times that are likely too short for the participant to attend to and read. **c**) Online error detection performance (n=254 trials). The inner circle shows proportions of correctly and incorrectly decoded words for all attempted words, along with skipped words which were displayed shorter than 300ms. The outer circle breaks down all displayed words based on both whether or not the correct word was decoded, and by whether neural error detection was successful or not after that word was presented.

Crucially, in this new task, a neural error classifier provided online feedback about whether it estimated that previously displayed words were correct or not based on the participant’s (neural) reactions. Specifically, an SVM classifier predicted the correctness of each decoded word based on the neural activity between 300 ms and 550 ms after that word was displayed. After the participant finished the whole sentence, the text of the decoded sentence was then colored based on the prediction of the error detector (**Fig. 5b**). During validation blocks, across 254 decoded words, 79.5% were correctly classified with 76.5% of errors detected at a false positive rate of 8.5% (**Fig. 5c**).

## Discussion

This study provides direct evidence that the human speech motor cortex encodes correlates of error processing during speech tasks. Using intracortical recordings from individuals with severe dysarthria, we identified temporally precise, feedback-locked neural responses encoding the correctness of word feedback. These responses were sufficiently robust to enable real-time decoding of speech BCI errors on a single-trial basis. From neural data alone, we could not only identify if an error had occurred, but also the category of the error.

These findings expand the known set of error-responsive cortical areas to include vPCG, an area traditionally associated with articulatory motor control. While feedback-related neural signals have been extensively documented in medial frontal cortex^6,8,7,9^ and premotor areas during manual motor tasks^18–20^, their presence in the speech motor cortex suggests that internal performance monitoring is a domain-general feature of motor systems, extending beyond limb movements to other complex, temporally structured behaviors like speech. Moreover, these findings complement traditional models, emphasizing a distributed and decentralized model of error-related feedback in the human brain in which task-relevant motor areas receive signals related to monitoring their own output.

Importantly, we also observed that neural activity in the speech motor cortex and IFG encoded not only whether an error was present—at a similar latency to the P300 component observed in EEG studies of surprise and outcome evaluation^7,34^—but also the nature of the error. These signals differentiated among phonological, orthographic, and semantic discrepancies between the intended and decoded words. This expands on previous descriptions of the N400 signals found in semantic conflict^35,36^, which emerges around 400 ms after the presentation of semantically incongruent stimuli. The timing and tuning of our intracortical signals are notably consistent with these findings, though our data capture intracortical neuronal-level responses rather than highly-attenuated scalp field potentials. Whether the neural encoding we observed represents a feedforward component of prediction error and/or feedback from linguistic circuits remains an open question.

These internally generated neural error signals open new opportunities for real-time, closed-loop control of speech BCIs. We demonstrated a proof-of-concept error detection system that successfully identified when the participant observed mistakes during a speech neuroprosthesis task paradigm that resembles naturalistic sentence production. This error decoder operated in parallel with a speech brain-to-text decoder that decoded the neural correlates of phoneme articulation^27,28^; thus, this combined system represents a hybrid “motor+cognitive” neuroprosthesis. These results suggest that neural error monitoring can be harnessed to enable corrective strategies, such as automatic word deletion or re-initiation of word/sentence decoding, without requiring explicit behavioral input from the user. This capability is particularly valuable in clinical populations with limited communication capacity, and it addresses a key challenge in BCI usability: reducing users’ burden of correcting system errors. Detecting error signals may also be useful for other purposes, for example as a biofeedback tool to help sharpen sensorimotor performance^37^.

Several limitations of this work warrant consideration. First, our paradigm focused primarily on visual feedback, which is relevant for BCIs designed to decode text. However, written text is not the natural modality for speech-related error monitoring. In naturalistic communications, speakers rely heavily on auditory feedback, including features such as pitch and formant dynamics. Paradigms that perturb these acoustic features have been instrumental in identifying rapid, trial-by-trial motor adaptations and associated cortical responses^23,38^. Replicating such paradigms in intracortical studies of speech error processing remains an important future goal. However, the modest fidelity of current voice neuroprostheses^26,39,40^ limits our ability to deliver real-time, intelligible auditory perturbations in closed-loop BCIs. More naturalistic feedback manipulations—e.g., introducing systematic mismatches in synthesized speech formants—could enable finer dissection of the neural dynamics underlying prediction error and adaptive control, particularly at the single-neuron level. Second, our sample size was limited to two individuals, each with distinct cortical array placements and both with vocal tract dysfunction due to ALS. While both exhibited similar error-related neural patterns, generalizing these findings to other populations or cortical recording locations will require broader validation. Third, while our analyses revealed that neural activity reflects categories of feedback mismatch, finer-grained investigations into how phonemic and semantic differences are encoded—such as systematically sampling along continua from minimal pairs to semantically related words—remain limited by the discrete and structured nature of the current task design.

## Methods

### Participants and ethics approvals

This study includes data from two participants, referred to as ‘T15’ and ‘T12’, who gave informed consent and were enrolled in the BrainGate2 clinical trial (ClinicalTrials.gov: NCT00912041). Permission for the clinical trial was granted by the U.S. Food and Drug Administration under an Investigational Device Exemption (Caution: Investigational device. Limited by U.S. federal law to investigational use). This manuscript does not report any primary clinical-trial related outcomes; instead, it describes scientific discoveries that were made using the data collected in the context of the ongoing clinical trial.

At the time of enrolling in the trial, T15 was a 45-year old left-handed man with ALS. He had tetraplegia and severe dysarthria, retained eye and neck movements, and retained limited orofacial muscle control. He had four 64-electrode Utah arrays (1.5 mm electrode length; Blackrock Neurotech, Salt Lake City, Utah) neurosurgically placed in his left vPCG; one in area 55b, two in area 6v and one in area 4 (Fig. 1B). See^28^ for more details. T12 was a 67-year old left-handed woman with ALS at the time of enrollment. She had dysarthria but retained partial use of her limbs. She had two 64-electrode Utah arrays placed in her left vPCG and two in area 44 (Broca’s area). See^27^ for more details. For both participants, the array location targets were identified through preoperative MRI scans and alignment of their brains to the Human Connectome Project^29^ cortical parcellation. Both participants could partially articulate and vocalize, but were unable to produce speech that was intelligible to an untrained listener.

### Neural signal processing and feature extraction

Raw neural signals were sampled at 30 kHz and preprocessed using a neural data acquisition system (NeuroPort Neural Signal Processor, Blackrock Neurotech) which bandpass filtered (0.3 Hz – 7.5 kHz, 4^th^ order Butterworth filter) and digitized the data into 1 ms data packets. The digitized packets were then received by our downstream data processing and decoding platform implemented using the BRAND framework^41^. The signals were further bandpass filtered (4th order zero-phase non-causal Butterworth filter) between 250 and 5000 Hz and then underwent array-wise Linear Regression Referencing^42,27,28^ to reduce noise artifacts. Putative neuronal action potentials (“threshold crossings”) and spike band power were computed using a 1 ms window per electrode. Threshold crossings were detected with a threshold at -4.5 times the root mean square voltage and a spike was registered if the voltage went below this threshold anytime within the window. Spike band power was computed as the square mean of the values within the same 1 ms window.

The scientific focus of this study is on whether task outcome error-related information is present in human speech cortical areas. The translational focus is on improving speech neuroprostheses; both of these questions are primarily concerned with ensemble-level neural codes (although we also show example single electrode multi-unit firing rates). Thus, rather than spike-sorting (isolating action potentials that are unambiguously from one neuron), we instead analyzed threshold crossings and spike band power. This choice substantially increases the yield of how many chronic array electrodes provide task-relevant neural signals, which is particularly important in human studies where re-implantation or electrode movement to improve neuronal isolation is not feasible^43^. We provide empirical and theoretical justifications in Trautmann et al., 2019 for why combining one or more neurons per electrode (as can be the case with threshold crossings and is the case for spike band power) does not preclude – and often facilitates – estimation of the underlying neural state compared to spike sorting.

### Eye tracking

T15’s gaze data were tracked using a Tobii Pro Spark eye tracker (Tobii AB, Stockholm, Sweden). During data collection blocks, T15’s on-screen gaze location was recorded at 60 Hz and used by the speech neuroprosthesis to allow him to select on-screen “buttons” by looking at them for 0.5 seconds.

### Behavioral tasks

All tasks employed an instructed-delay paradigm. Each trial began with the visual presentation of a cue and a “delay” period, indicated by a red square, during which the participant read and prepared to act on the cue. This was followed by a “go” cue (the square turning green), prompting the participant to initiate their response. The duration and structure of the go period varied by task.

In the closed-loop BCI tasks—including the Sentence Copy, Sequential Feedback, and Emphasized Feedback tasks—speech attempts were self-paced. The participant ended each trial by fixating on an on-screen “button” using an eye tracker-controlled cursor. In contrast, the open-loop Word Copy Task featured a fixed 2-second window for speech attempts following go cue presentation.

Neural recordings were collected with participant T15 during the BCI Sentence Copy, Sequential Feedback, Emphasized Feedback, and Word Copy tasks. With participant T12, only the Word Copy Task was performed.

### BCI Sentence Copy Task

In the Sentence Copy Task, a sentence prompt was displayed on the screen during the 4-second delay period. After the go cue, participant T15 attempted to speak the full sentence. A phoneme decoder and a 5-gram language model processed the neural data continuously, generating predictions every 80 ms. The predicted sentence was dynamically updated and displayed on the screen in real time. Once the participant completed the utterance, he ended the trial using the eye tracker interface.

Following sentence completion, a large language model re-ranked candidate sentences, and the top prediction was rendered into synthesized voice using a personalized text-to-speech (TTS) synthesizer modeled on the participant’s pre-ALS voice. For full system architecture and decoding methods, see Card et al. (2024).

### Sequential Feedback Task

In this task, the participant was shown a short sentence (fewer than five words) during the delay period and instructed to memorize it, as the sentence would no longer be visible during the go period. This design was intended to simulate natural speech planning rather than reading out loud.

During the go phase, the participant attempted to speak one word at a time. A phoneme decoder and language model continuously predicted phoneme probabilities and word candidates. If two consecutive “silence” phoneme logits were detected, the system inferred the end of a word attempt and aligned the latest language model prediction with the sentence prompt using a dynamic edit distance algorithm. The task then identified the word being attempted, and displayed the most recently predicted word on the screen. Upon trial completion, triggered via eye tracker, the sequence of previously displayed feedback words was assembled as the final decoded sentence.

### Emphasized Feedback Task

This task followed the same basic structure as the Sequential Feedback Task, with the participant memorizing a short sentence and speaking it word by word after the go cue. However, it introduced two key modifications. The first was intended to help direct the participant’s attention upon the feedback word right when it was displayed: the latest decoded word was visually highlighted (centered in a separate row with a blue background and a larger font) immediately upon prediction. The second modification was a real-time demonstration of an error detection module that operated in parallel to the word decoder.

The highlight faded after 600 ms or upon the prediction of the next word. A closed-loop SVM-based error decoder used neural activity during the feedback window (300–450 ms post-word display) to estimate whether the displayed word matched the participant’s intended utterance. Words shown for less than 300 ms were excluded from error decoding (“skipped”), as they likely did not afford sufficient visual attention. At the end of the trial, the full decoded sentence was displayed with predicted errors highlighted in red, skipped words in gray, and correctly decoded words in white.

### Word Copy Task

In the Word Copy Task, participants were presented with a single word during the delay period. After the go cue (delivered 2 seconds later), the cue word disappeared from the screen and the participant attempted to articulate it. For participant T12, the word attempt duration was fixed at 2 seconds; for participant T15, the trial ended automatically upon neural phoneme decoding detection of sustained silence.

Following the speech attempt, a feedback word was displayed. Each cue word was associated with one of five possible feedback conditions, which were designed to vary in their phonological and semantic similarity to the original cue. These conditions were: 1) the cue word itself (correct feedback); 2) a synonym that was semantically similar to the cue word; 3) a minimal-pair word that differed by a single phoneme; 4) a pseudoword that was not a real English word but differed from the cue word by one phoneme at the same position as the minimal pair; and 5) a homophone that shared the same phoneme sequence as the cue word but had different spelling and meaning. Phonological differences were computed using the ‘g2p-en’^45^ grapheme-to-phoneme conversion tool, and semantic similarity was quantified using cosine-similarity between BERT-based^46^ embeddings. The complete set of cue-feedback pairs is provided in **Table 1**. During the task, the frequency of each feedback condition was roughly 3:1:1:1:1 (cue:pseudoword:homophone:minimal pair:synonym) to maintain the sense that this task was similar to (somewhat inaccurate) BCI decoding for the participants.

To increase participant engagement and confirm his attention to the feedback, participant T15 was asked to indicate whether the feedback word matched the cue using eye-tracked saccades. The direction-to-response mapping was counterbalanced across trials: for half of the blocks, the participant was told to use leftward gaze to indicate that the feedback was correct and rightward gaze to indicate it was incorrect; for the other half, this mapping was reversed. This design allowed us to dissociate error-related neural activity from the direction of the eye movement itself.

To investigate error encoding in response to auditory feedback, we implemented a variant of the Word Copy Task in which, following participant T15’s speech attempt, feedback was delivered as synthesized speech rather than as a visual word. The spoken feedback was generated using the same personalized TTS model described in the BCI Sentence Copy Task and was played aloud through a speaker.

### Data analyses

#### Single electrode modulation analyses

To assess how neural activity measured at individual electrodes modulated based on feedback correctness (Fig. 1d,e, Fig. 2c, Fig. 4b, Supplemental Fig. 4b) and error type (Fig. 3a), we analyzed threshold crossings binned at 10 ms intervals from -200 to 1000 ms relative to feedback onset. The data were smoothed with a one-dimensional Gaussian kernel (σ = 20 ms) and trial-averaged. To evaluate the electrodes that showed significant tuning to word correctness, we computed spike band power in the 200–600 ms window post-feedback and compared responses to correct vs. incorrect feedback using a Wilcoxon rank-sum test (Supplemental Fig. 2a).

#### Population modulation analyses

To quantify the population-level neural modulation in response to feedback, we used a cross-validated neural distance metric^47^. Specifically, we computed the squared distance between neural responses (based on binned spike band power) to feedback versus either resting baseline activity or alternate feedback conditions (e.g., correct vs. incorrect feedback; Fig. 2d, Supplemental Fig. 2d,h; or across error types; Fig. 3b, Supplemental Fig. 3a,d). The neural distance for each condition was calculated as:

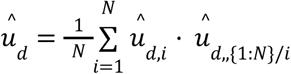

Here, 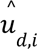 is the estimate of the difference in means using single trial *i* from both distributions and 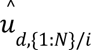 is the estimate of the difference in means using all trials except trial *i* from the distributions. For imbalanced datasets, K-fold cross-validation was used, with the number of splits determined by the smaller class size. To assess temporal dynamics, we computed neural distance using a 50 ms sliding window with a 10 ms step.

#### Gaze analysis

To infer which feedback word was being viewed in the BCI Sentence Copy Task, we defined fixations as gaze points remaining within a 100-pixel radius of the screen facing the participant for longer than 300 ms. Fixations separated by less than 50 ms were merged to account for blinks and eye tracker noise. In the Word Copy Task, saccade onset, which was used as a behavioral response, was defined as a gaze shift exceeding 15 pixels/sec and 200 pixels from the screen center.

#### Offline error decoder training and analyses

To decode whether the word feedback provided to the participant was correct or incorrect based on their neural response, or to classify the type of feedback error, we trained SVM classifiers using spike band power features. Unless otherwise noted, T15’s d6v array was excluded from decoding due to its minimal contribution of error-related signals.

For the Word Copy Task, trials were labeled as incorrect if the feedback word differed from the cue word. To avoid overfitting to word identity, we applied leave-one-cue-out cross-validation: all trials corresponding to one of the six cue words were held out in each fold. Within each fold, we used random oversampling to balance class sizes, and averaged classification performance across folds. This cross-validation procedure was repeated 10 times to account for variability due to resampling. Chance-level performance was estimated using 1000 iterations of label shuffling. Classification was performed on 150 ms sliding windows with 10 ms stride within a window spanning [-200, 1000] ms around feedback onset.

To better characterize when the neural ensemble response corresponding to a specific type of error feedback can be best differentiated from that of the other error types, we combined the confusion matrix from different time points by taking highest recall rates that are significantly above chance for each row (**Supplemental Fig 4c,f1**). Statistical significance was assessed by comparing classification performance to a null distribution obtained from 1000 label permutations.

In the BCI Sentence Copy and Sequential Feedback tasks, determining feedback word correctness first required determining what word in the promoted sentence each decoded word corresponded to. We aligned the decoded and prompted sentence using an edit-distance-based word matcher. For time points where multiple words were located within the radius centered on the participant’s gaze fixation, all of these words needed to be correct to assign a “correct” label. We used stratified 5-fold cross-validation, with oversampling and label shuffling procedures as described above. For these analyses, a 100 ms sliding window (10 ms stride) was used for classification.

#### Online error decoder training and inference

To enable real-time error detection in the Emphasized Feedback Task, we first pretrained an SVM classifier on data from the Sequential Feedback Task. During online use, binned spike band power features were collected between 300 and 550 ms after each decoded word’s on-screen feedback onset and processed using overlapping 100 ms windows (50 ms stride). The median of the resulting logits was compared to a fixed threshold to determine whether the most recently displayed word was classified as correct or incorrect. This architecture accommodated variability in reaction times to when decoded words appeared on-screen.

## Data and code availability

De-identified data and the codes used in these analyses will be available upon publication.

## Acknowledgements

We thank participants T15 and T12, their families and care partners for their contributions to this research.

This work is supported by the A.P. Giannini Foundation Postdoctoral Research Fellowship (Card); the Office of the Assistant Secretary of Defense for Health Affairs through the Amyotrophic Lateral Sclerosis Research Program under award number AL220043; DP2 from the NIH Office of the Director and managed by NIDCD (1DP2DC021055); Searle Scholars Program; and a Career Award at the Scientific Interface from the Burroughs Wellcome Fund (Stavisky).

## Author contributions

X.H. led the experiments, analyzed the data, created the figures and implemented error decoder on the real-time data acquisition system. X.H. and C.I. interfaced with participant T15, scheduled research sessions and collected the primary data for this study. N.S.C., M.W., X.H. and T.S.C. developed T15’s brain-to-text BCI. X.H., E.M.K., C.F., F.K., N.H. interfaced with participant T12 and collected data for this study. J.M.H. and F.R.W. supervised research at the Stanford site. L.R.H. is the sponsor-investigator of the multisite clinical trial. D.M.B. was responsible for all clinical trial-related activities at U.C. Davis. X.H., S.D.S., and D.M.B. were involved in conceptualization of the study and experimental design. S.D.S. and D.M.B. supervised all aspects of the project. X.H., S.D.S., and D.M.B. wrote the manuscript. All authors reviewed and helped edit the manuscript.

## Competing interests

IDE Caution Statement: CAUTION: Investigational Device. Limited by Federal Law to Investigational Use. The content is solely the responsibility of the authors and does not necessarily represent the official views of the National Institutes of Health, or the Department of Veterans Affairs, or the United States Government. The MGH Translational Research Center has a clinical research support agreement (CRSA) with Ability Neuro, Axoft, Neuralink, Neurobionics, Paradromics, Precision Neuro, Synchron, and Reach Neuro, for which L.R.H provides consultative input. L.R.H is a non-compensated member of the Board of Directors of a nonprofit assistive communication device technology foundation (Speak Your Mind Foundation). Mass General Brigham (MGB) is convening the Implantable Brain-Computer Interface Collaborative Community (iBCI-CC); charitable gift agreements to MGB, including those received to date from Paradromics, Synchron, Precision Neuro, Neuralink, and Blackrock Neurotech, support the iBCI-CC, for which L.R.H provides effort. M.W., S.D.S. and D.M.B. have patent applications related to speech BCI owned by the Regents of the University of California including IP which has been licensed to a neurotechnology startup. SDS, JMH and FRW are inventors on intellectual property licensed by Stanford University to Blackrock Neurotech and Neuralink Corp. JMH is a consultant for Paradromics, serves on the Medical Advisory Board of Enspire DBS and is a shareholder in Maplight Therapeutics. He is also the co-founder of Re-EmergeDBS. S.D.S. is an advisor to Sonera. D.M.B. is an ad-hoc consultant for Globus Medical Inc., and was an ad-hoc consultant for Paradromics Inc. during the period of data collection for this manuscript.

## Supplemental Figures

**Supplemental table 1.** Data collection sessions.

| Session | Tasks |
| --- | --- |
| T12.2024.01.11 | Word Copy Task (Supplemental Fig. 2 & 3) |
| T15.2024.02.16 | Word Copy Task (Fig. 2 & 3, Supplemental Fig. 2 & 3) |
| T15.2024.07.21 | Brain-to-text Task (Fig. 4) |
| T15.2024.08.04 | Sequential Feedback Task (Fig. 1) |
| T15.2024.08.23 | Emphasized Feedback Task (Fig. 5) |
| T15.2024.11.24 | Word Copy Task w/ auditory feedback (Supplemental Fig. 4) |

**Supplemental Fig. 1.**
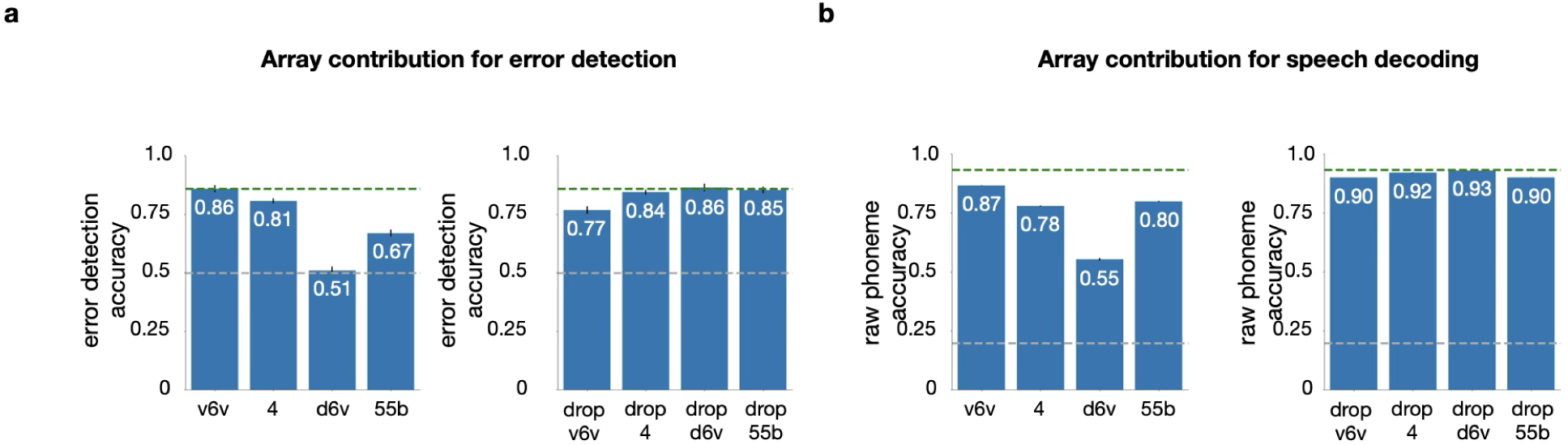
Array contribution for error detection versus speech decoding. **a**) Offline classification accuracy of the correctness of the feedback (mean ± s.d.) for individual arrays (left) and without each array (right) at the optimal classification window indicated in Fig.1. These data come from the brain-to-text task with sequential feedback. **b**) Raw phoneme error rates derived from the speech decoder output during a standard brain-to-text task^28^, broken down by individual arrays. *Left*: decoding performance while using each array alone. *Right:* performance when excluding one array at a time. The green dashed lines show classification accuracy using all four arrays while grey dashed lines show chance-level performance. Taken together these results indicate that the relative importance of arrays was similar for both speech decoding and error detection.

**Supplemental Fig. 2.**
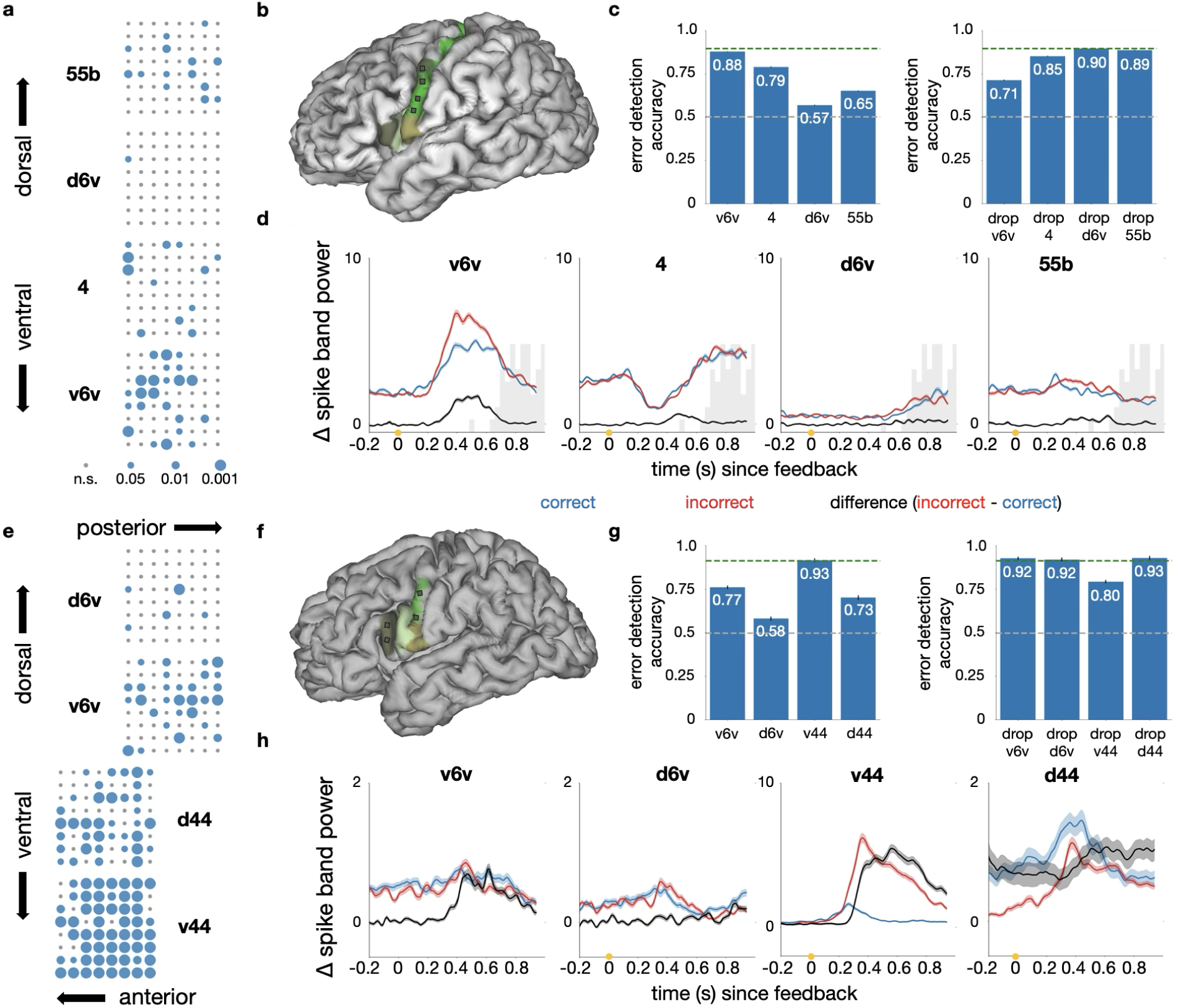
Neural error signal during a Word Copy Task in two participants. **a**), **e**) Electrode array maps showing electrodes with significant tuning for error feedback. Significance was determined using Wilcoxon two-sided rank sum test comparing neural responses after incorrect and correct feedback. Blue dots indicate significant electrodes, with dot size corresponding to p values. **b**)&**f**) Approximate array locations (grey squares) superimposed on a three dimensional reconstructions of participant T15’s and T12’s brains, respectively. Colored regions correspond to cortical areas aligned to the participant’s brain with the use of the Human Connectome Project MRI protocol scans before implantation. **c**)&**g**) Offline classification accuracy (mean ± s.d.) of feedback correctness at 410 ms post-feedback. Left, accuracy for individual arrays. Right, accuracy when arrays in specific brain areas are excluded or exclusively used. Green dashed lines indicate performance using all arrays, while gray lines represent chance-level accuracy. **d**)&**h**) Neural population modulation (mean ± s.e.) in spike band power for individual arrays of each participant. Blue and red lines indicate neural distances between feedback (correct and incorrect, respectively) responses and a “do nothing” baseline condition. The black line reflects the neural distance between incorrect and correct feedback responses. Note the different vertical scales. The light gray histogram in panel d indicates the time distribution of when participant T15 initiated a response with a saccade.

**Supplemental Fig. 3.**
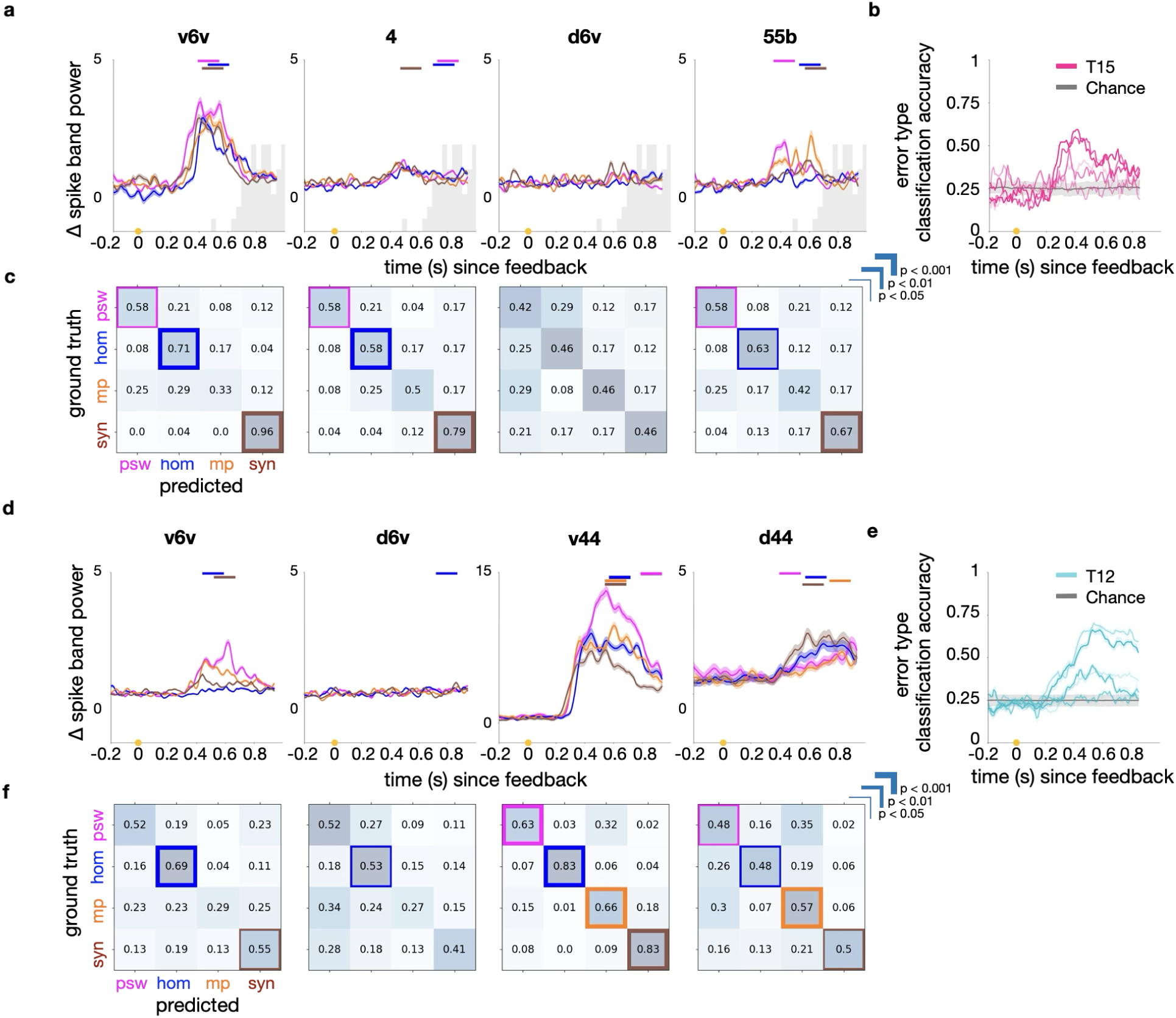
Error type classification in both participants. **a**)&**d**) Neural population modulation (mean ± s.e.) for each incorrect feedback type in the spike band power, presented for participants T15 (top) and T12 (bottom). Modulation is quantified as the neural distance between each incorrect feedback type and the correct feedback type (note that this doesn’t necessarily capture differences *between* error types). Colored lines above the plots indicate time windows when each feedback type could be most accurately identified (versus the other types), i.e., optimal recall time windows. Note the different vertical scale for T12’s v44 array. The light gray histogram in panel a indicates the time distribution of when participant T15 initiated a response with a saccade. **b**)&**e**) offline classification accuracy of incorrect feedback types using neural features from each array. The two more accurate arrays for T15 were the v6v and 4 arrays. For T12, it was the v44 and d44 arrays. **c**)&**f**) Confusion matrices for feedback type classification, optimized for the best recall time window for each row. Rows represent individual feedback types, and significant diagonal cells (Bonferroni-corrected) are highlighted with colored boxes of varying thickness. The classification windows for each feedback type, corresponding to their optimal time points, are marked with matching colored lines in panels (a, d).

**Supplemental Fig. 4.**
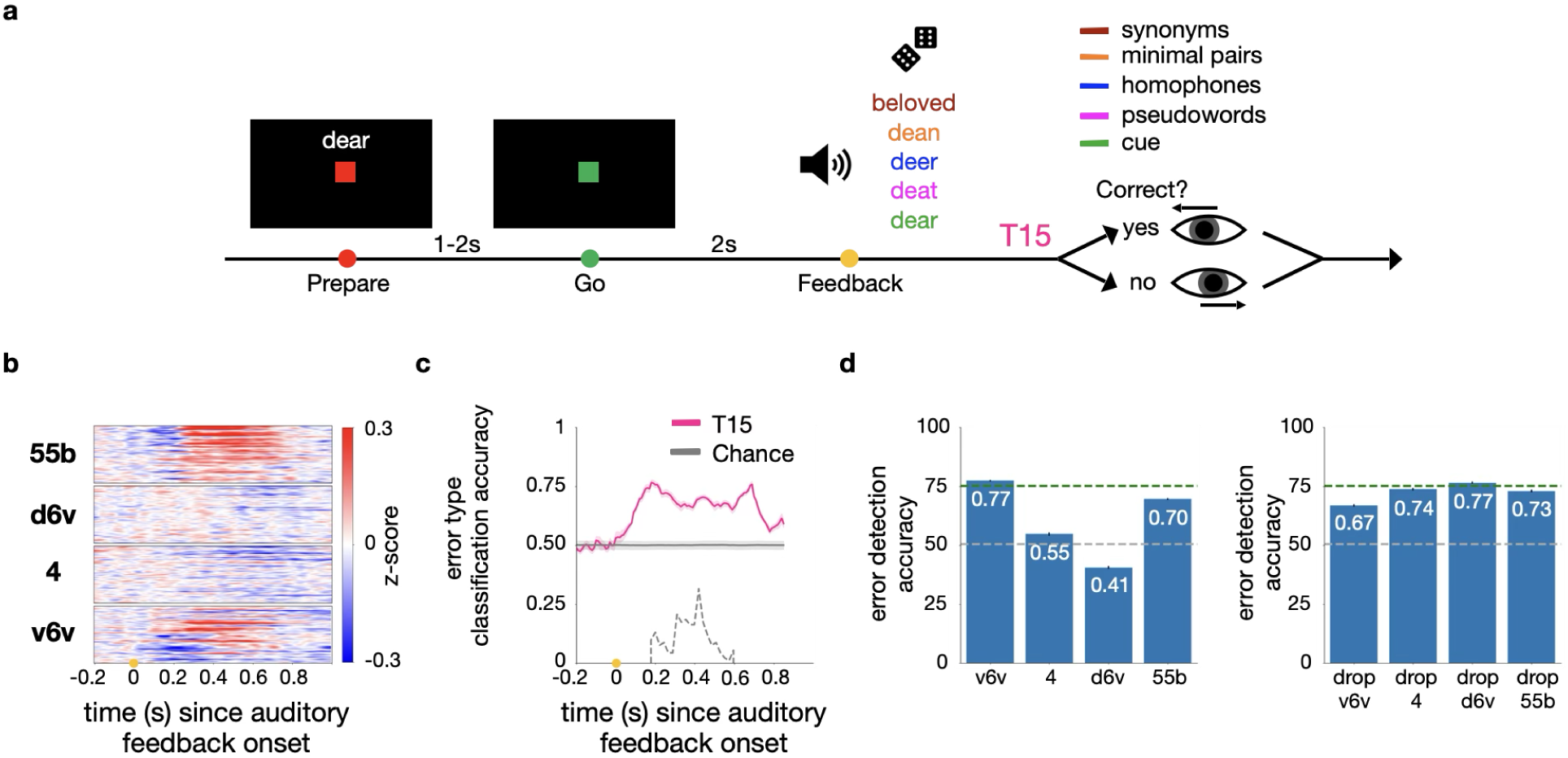
Neural error signals during Word Copy Task with auditory instead of visual feedback. **a**) Illustration of the Word Copy Task variant with auditory feedback. Participant T15 was prompted to say a single word. After he attempted to speak it, synthesized speech from a text-to-speech (TTS) model corresponding to one of five feedback types (the cue word, a pseudoword, a homophone, a minimal pair, or a synonym) was played through a speaker. He was prompted to respond to confirm feedback correctness using an eye tracker. **b**) Normalized trial-averaged spike band power difference between incorrect and correct feedback conditions. Neural activity is aligned to the onset of the TTS-synthesized speech (n=209 trials; 88 correct and 121 incorrect). **c**) Offline classification accuracy of the correctness of the feedback (mean ± s.d.) as a function of the start of a 150-ms sliding window of neural data. The light gray dashed line indicates the time distribution of when audio feedback ended. **d**) Offline classification accuracy (mean ± s.d.) of feedback correctness at 180 ms after playback onset. *Left:* decoding performance while using each individual array. *Right:* performance when excluding one array at a time. The green dashed lines represent classification accuracy using all four arrays while grey dashed lines represent chance-level performance.

